# Spastin is an essential regulator of male meiosis, acrosome formation, manchette structure and nuclear integrity

**DOI:** 10.1101/2022.08.01.502419

**Authors:** Samuel R. Cheers, Anne E. O’Connor, Travis K. Johnson, D. Jo Merriner, Moira K. O’Bryan, Jessica E. M. Dunleavy

**Affiliations:** School of BioSciences and Bio21 Institute, The University of Melbourne, Parkville, VIC 3010, Australia; School of Biological Sciences, Monash University, Clayton, VIC 3800, Australia

**Author notes:** Equal senior authors (Moira K. O’Bryan, Jessica E. M. Dunleavy).

**Keywords:** male infertility, spermatogenesis, microtubule severing, AAA ATPase, hereditary spastic paraplegia

## Abstract

The development and function of male gametes is critically dependent on a dynamic microtubule network, yet how this is regulated remains poorly understood. We have recently shown that microtubule severing, via the action of the meiotic AAA ATPase protein clade, plays a critical role in this process. Here, we sought to elucidate the roles of spastin, an as yet unexplored member of this clade in spermatogenesis. Using a *Spast*^*KO/KO*^ mouse model, we reveal that spastin loss resulted in a complete loss of functional germ cells. Spastin plays a critical role in the assembly and function of the male meiotic spindle, and in its absence, apoptosis is significantly increased. Consistent with meiotic failure, round spermatid nuclei were enlarged, indicating aneuploidy, but were still able to enter spermiogenesis. During spermiogenesis, we observed extreme abnormalities in manchette structure, supernumerary acrosome formation, and commonly, a loss of nuclear integrity. This work defines a novel and essential role for spastin in regulating microtubule dynamics during spermatogenesis and is of potential relevance to patients carrying Spastin variants and to the medically assisted reproductive technology industry.

**Summary statement:** We identify an essential role for the microtubule severing enzyme spastin in the regulation of microtubule dynamics during spermatogenesis.

## Introduction

Microtubule severing is fundamental to the regulation of microtubule dynamics and is achieved via members of the meiotic clade of the AAA superfamily (‘ATPases associated with diverse cellular activities’). This group includes the katanins, the fidgetins, and spastin (SPAST), which all have microtubule severing activity, in addition to VPS4, which has no known severing function (Frickey and Lupas, 2004). While the utility of microtubule severing in most mammalian developmental processes is unexplored, critical roles in neurodevelopment are well established for spastin and the katanins (Ahmad et al., 1999, Chen et al., 2014, Hu et al., 2014, Tan et al., 2020, Yu et al., 2008, Liu et al., 2021), and in multiple aspects of male germ cell development, for the katanins (Dunleavy et al., 2021, Dunleavy et al., 2017, O’Donnell et al., 2012). *SPAST* mutations are the most common cause of hereditary spastic paraplegia (Hazan et al., 1999). Hereditary spastic paraplegia is caused by progressive degeneration of neurons in the central nervous system and is characterised by lower limb stiffness, weakness, and spasticity. *SPAST* mutation is dominant, however, it is still debated whether disease is caused by haploinsufficiency or by gain-of-function, though it appears an interplay of both mechanisms is likely (Qiang et al., 2019). Although spastin is expressed in other tissues, its roles outside the nervous system remain virtually unexplored. Of direct relevance to this study, spastin is highly expressed in male germ cells.

As is typical of most members of the meiotic AAA clade, spastin is defined by a highly conserved AAA domain, an N-terminal microtubule interacting and trafficking (MIT) domain, and a C-terminal VPS4 domain that has been implicated in oligomerisation (Snider et al., 2008, Vajjhala et al., 2008, Rigden et al., 2009). The AAA domain confers the ATPase activity necessary for microtubule severing (Erdmann et al., 1991). Outside the AAA domain, the MIT domain binds microtubules to increase severing efficiency (Errico et al., 2002, Roll-Mecak and Vale, 2008) and is used for interactions with components of the ESCRT-III (endosomal sorting complexes required for transport) machinery (Reid et al., 2005, Yang et al., 2008). Additionally, spastin contains a microtubule-binding domain (MTBD) that is essential for microtubule severing (White et al., 2007). A unique feature of spastin among the meiotic AAA clade is the presence of a hydrophobic region at the N-terminal that allows it to embed within lipid membranes. A second smaller isoform of spastin is ubiquitously expressed that lacks this N-terminal hydrophobic region and is found throughout the cytosol of COS7 cells (Claudiani et al., 2005).

To sever microtubules, ATP-bound spastin subunits assemble into a spiral-shaped homohexamer around the C-terminal tail of tubulin (Roll-Mecak and Vale, 2008, Sandate et al., 2019). Upon hydrolysis of ATP, the hexamer changes conformation to a ring and in this movement tugs upon the tubulin C-terminal tail, removing the tubulin heterodimer from the microtubule lattice (Roll-Mecak and Vale, 2008, Zehr et al., 2020). The action of spastin and other microtubule-severing enzymes can lead to microtubule disassembly, the release of microtubules from nucleation sites, and the generation of short stable seeds of microtubules for transport to other parts of the cell and/or to nucleate microtubule growth (reviewed in (McNally and Roll-Mecak, 2018)). Conversely, and perhaps counterintuitively, spastin action can lead to microtubule stabilisation by removing GDP-associated tubulin heterodimers from the microtubule lattice, which are then replaced with more stable GTP-associated tubulin heterodimers (Vemu et al., 2018).

Cellular functions for spastin include the severing of microtubules at the spindle poles in *D. melanogaster* during mitosis to allow poleward movement of chromosomes (Zhang et al., 2007), shaping of the endoplasmic reticulum in cultured rat neurons, HEK293 and COS7 cells (Park et al., 2010), development of the axon through microtubule outgrowth in zebrafish neurons (Wood et al., 2006), and axonal transport in isolated squid axoplasm (Leo et al., 2017). To date, studies conducted using rodent models have focused solely on brain development and have identified a role for spastin in neurogenesis, axonal development, and axonal transport (Ji et al., 2018, Kasher et al., 2009, Jeong et al., 2019).

Through the MIT domain, spastin is able to interact with components of the ESCRT-III machinery (Reid et al., 2005, Yang et al., 2008). Through these interactions, spastin is involved in endosome formation and processing, nuclear envelope reformation, and midbody abscission during cytokinesis (reviewed in (Migliano et al., 2022)). ESCRT-III recruits spastin to the midbody during mitotic cytokinesis in HeLa cells to sever the midbody microtubules and allow the completion of membrane fission (Connell et al., 2009). Similarly, after cell division in HeLa cells, ESCRT-III recruits spastin to sites on the reforming nuclear membrane through which microtubules pass. Severing of these microtubules allows for the sealing of the nuclear membrane (Vietri et al., 2015). Spastin is also required in HeLa cells and mouse embryonic fibroblasts for endosomal tubulation and fission and correct lysosome function (Allison et al., 2013, Allison et al., 2017). Both of these functions require interaction with the ESCRT-III components and the ability of spastin to sever microtubules. Finally, spastin is involved in the movement and metabolism of lipid droplets in HeLa cells, which, interestingly, requires interaction with ESCRT-III components but not microtubule severing (Chang et al., 2019, Arribat et al., 2020).

The development of male germ cells, like that of neurogenesis, is highly dependent on complex microtubule structures. These include the bipolar spindle and midbody during mitosis and meiosis, the manchette for sperm head shaping, and the axoneme which forms the core architecture of the sperm tail. Previous research has shown that spermatogenesis is critically dependent on microtubule severing through other members of the meiotic group of AAA proteins, the katanins. KATNAL1 is required for regulation of microtubule dynamics within the Sertoli cells; the somatic support cells within the seminiferous epithelium of the testis (Smith et al., 2012). KATNAL2 is required for the suppression of supernumerary centriole formation and for sperm tail formation, sperm head shaping, and sperm release from the seminiferous epithelium (Dunleavy et al., 2017). Finally, loss of function of KATNB1, the regulatory katanin subunit, results in failures in meiosis, as well as in acrosome formation, sperm head shaping, and several aspects of tail formation (Dunleavy et al., 2021, O’Donnell et al., 2012). The role of spastin in the development of male germ cells has not yet been directly tested. However, spastin is highly expressed in the testis (Karlsson et al., 2021), and, notably, a previous publication reported that homozygous spastin mutant mice were male sterile, but the biological origin of this phenotype was not investigated (Tarrade et al., 2006).

Here, we directly tested the role of spastin in spermatogenesis using a whole-body *Spast* knockout mouse model in which a truncation occurred after exon 4. We reveal that spastin is essential for male germ cell development in the mouse and loss of spastin is incompatible with the production of male germ cells. Our work identifies spastin as a regulator of anaphase during meiosis, of acrosome biogenesis, and of the sculpting of the sperm head via the manchette. Interestingly, we also find that spastin plays a critical role in maintaining haploid germ cell nuclear integrity. Spastin loss leads to male sterility characterized by incomplete meiotic failure, followed by catastrophic degeneration of the spermatid structure.

## Results

### Spastin is required for spermatogenesis and male fertility

To investigate the role of spastin in male germ cell development, we used a whole-animal *Spast* knockout mouse model (*Spast*^*KO/KO*^) comprising a trans-NIH Knockout Mouse Project (KOMP) construct (Fig. 1A) inserted into *Spast* intron 4 (red arrowhead, Fig. 1B,C) designed to truncate *Spast* mRNA at exon 4. The homozygous presence of the construct resulted in an 89.5% reduction in *Spast* mRNA expression in *Spast*^*KO/KO*^ compared to *Spast*^*WT/WT*^ testes (Fig. 1D). Sequencing of the PCR product confirmed that low levels of *Spast* mRNA containing sequences from the construct were produced in the *Spast*^*KO/KO*^ mouse. This indicates that, in common with several other KOMP constructs, a low degree of transcription occurred. Due to the presence of the construct in the mRNA, it is unlikely that translation would result in functional spastin. This is the first study to use this construct as a whole-body knockout. Previous studies using this mouse model generated a tissue-specific conditional knockout (Magiera et al., 2018, Brill et al., 2016).

**Fig. 1:**
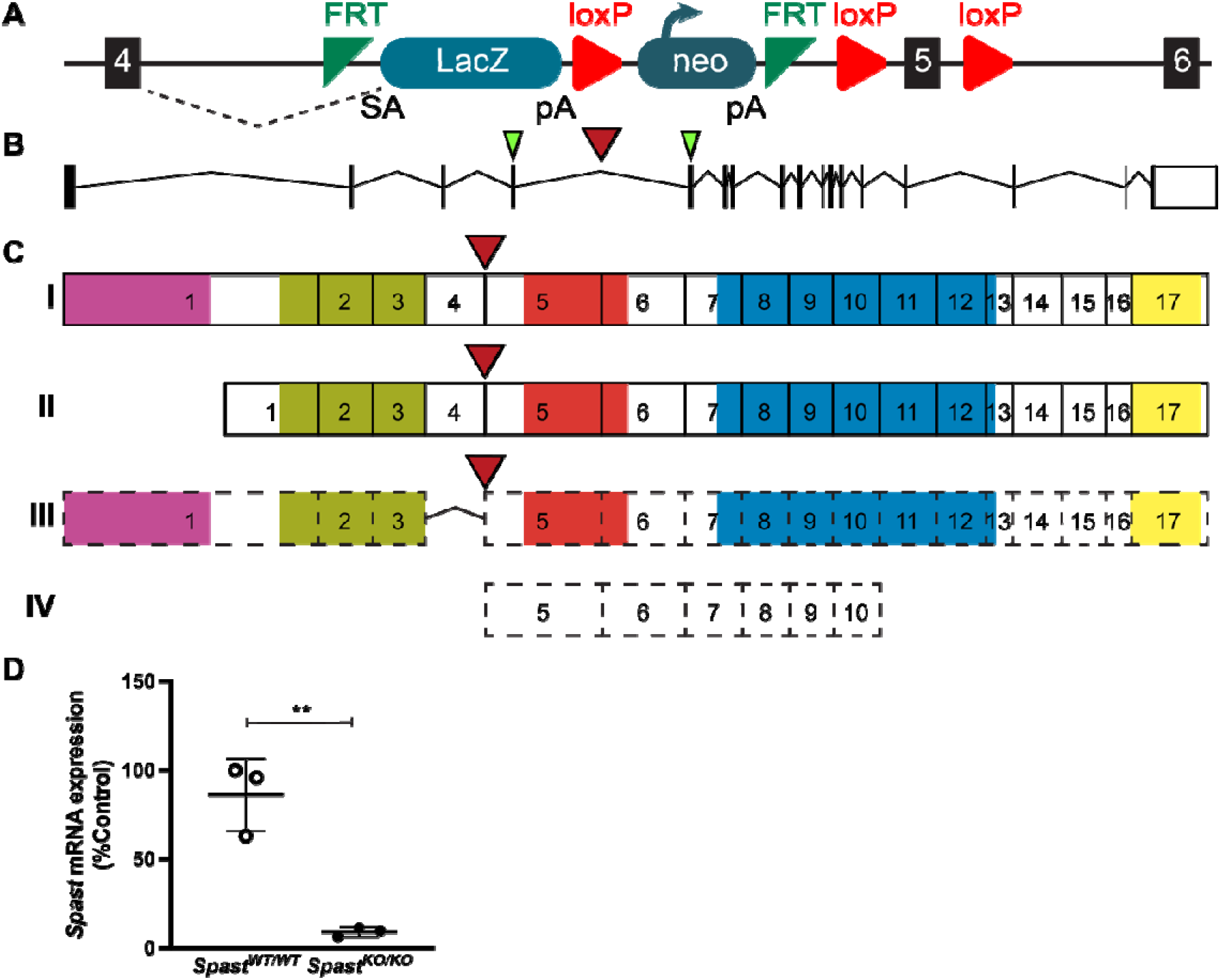
Ablation of spastin function in *Spast*^*KO/KO*^ mice. (**A**) The *Spast* KO-first conditional ready allele. The FRT-lacZ-loxP-neo-FRT-loxP-*Spast* exon 5-loxP cassette was inserted into *Spast* intron 4. (**B**) Schematics of the murine *Spast* gene and (**C**) SPAST protein. The red arrow indicates the point of cassette insertion. The green arrowheads indicate the target regions of the qPCR primers. The hydrophobic region, MIT domain, MTBD domain, AAA ATPase domain, and VPS4 oligomerisation domain are shown in pink, green, red, blue, and yellow, respectively. Two *Spast* isoforms M1 (**I**) and M87 (**II**) have been characterised in mice (Mancuso and Rugarli, 2008). Two additional *Spast* isoforms (**III, IV**), that are yet to be validated, are predicted (Uniprot, A0A286YCJ4 and A0A3B2WBA7). (**D**) qPCR analysis of *Spast* transcript levels in the *Spast*^*WT/WT*^ and *Spast*^*KO/KO*^ whole testes (n= 3 mice/genotype). Lines represent the mean ± SD and are normalised to the expression of *Ppia*. \*\**P* = 0.0028.

*Spast*^*KO/KO*^ mice generated from the intercrossing of heterozygous mice were born at the expected Mendelian frequency. *Spast*^*KO/KO*^ male mice exhibited normal mating behaviour when partnered with *Spast*^*WT/WT*^ female mice, however they were uniformly male sterile (8.5 pups per copulatory plug in *Spast*^*WT/WT*^ (n=3) versus 0.0 pups in *Spast*^*KO/KO*^ (n=4), *p* = <0.0001). Analysis of the *Spast*^*KO/KO*^ male reproductive tract revealed the complete absence of sperm. *Spast*^*KO/KO*^ mice had normal body weight, but significantly smaller adult testes compared to *Spast*^*WT/WT*^ controls (36.7% reduction; Fig. 2A). An analysis of testis daily sperm production revealed that *Spast*^*KO/KO*^ mice produced 99.4% fewer sperm (Fig. 2B), and their epididymal sperm content was reduced by 99.7%, compared to *Spast*^*WT/WT*^ controls (Fig. 2C). Rare cells that were seen in the epididymis of *Spast*^*KO/KO*^ mice were prematurely sloughed spermatocytes and round spermatids rather than spermatozoa (Fig. 2D,E, blue arrowheads). This was in stark contrast to *Spast*^*WT/WT*^ epididymides, which were full of spermatozoa (Fig. 2E, cauda).

**Fig. 2:**
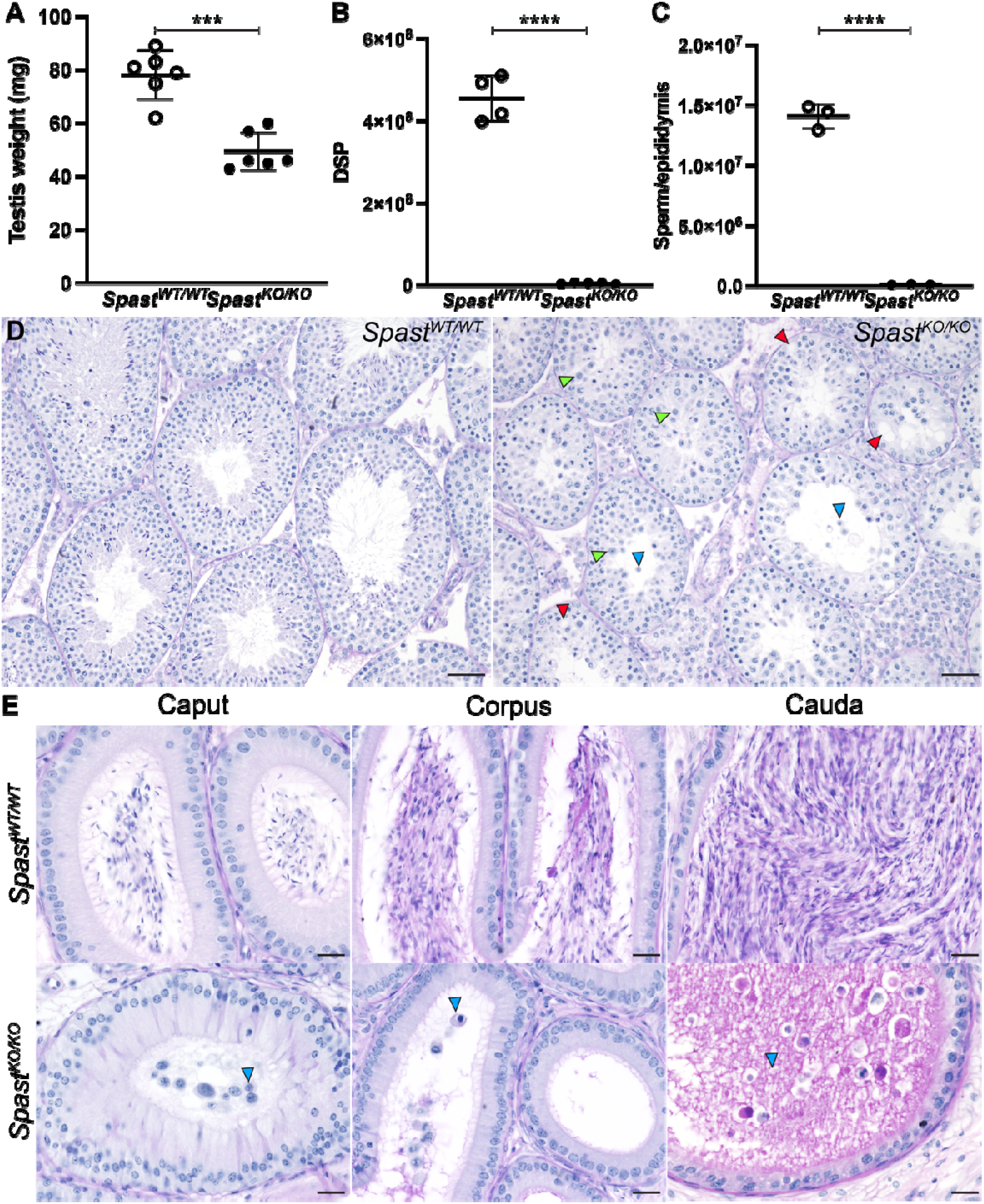
Spermatogenic defects due to knockout of *Spast*. (**A**) Testis weight, (**B**) total daily sperm production (DSP) per testis, and (**C**) epididymal sperm content in *Spast*^*KO/KO*^ mice (black circles) compared to *Spast*^*WT/WT*^ (white circles) controls (n≥3 mice/genotype, lines represent mean ± SD). Asterisks denote different levels of significance; *** p= 0.0001, **** p< 0.0001. (**D**) PAS-stained testis sections from *Spast*^*WT/WT*^ and *Spast*^*KO/KO*^ mice. Red arrowheads indicate vacuoles in the seminiferous epithelium. The green arrowheads indicate abnormally large round spermatids. Blue arrowheads indicate prematurely released germ cells. Scale bars = 50µm. (**E**) Epididymis sections from *Spast*^*WT/WT*^ and *Spast*^*KO/KO*^ mice. Blue arrowheads indicate prematurely released germ cells. Scale bars = 20 µm.

Histological analysis of *Spast*^*KO/KO*^ testes identified multiple abnormalities at various stages of spermatogenesis. Consistent with premature germ cell sloughing and/or death, large areas of the seminiferous epithelium were devoid of germ cells and/or exhibited a ‘lacy’ appearance in *Spast*^*KO/KO*^ testes, indicative of recent germ cell loss (Fig. 2D, red arrowheads). In the majority of *Spast*^*KO/KO*^ seminiferous tubules, spermatogonia and primary spermatocytes up to and including prophase I appeared phenotypically normal. The earliest point at which a clear defect could be seen in the *Spast*^*KO/KO*^ mice was during metaphase of meiosis I. During meiotic division, cells often displayed misaligned chromosomes and signs of division failure, resulting in abnormally large round spermatids (Fig. 2D, green arrowheads) or, rarely, bi-nucleated spermatids (Fig. 3B, orange arrowhead), suggesting that spastin may play a critical role in meiosis. We also noticed that round spermatids in *Spast*^*KO/KO*^ testes had abnormal acrosome development and failed to elongate, indicating an essential role for spastin during the early processes of spermiogenesis.

**Fig. 3:**
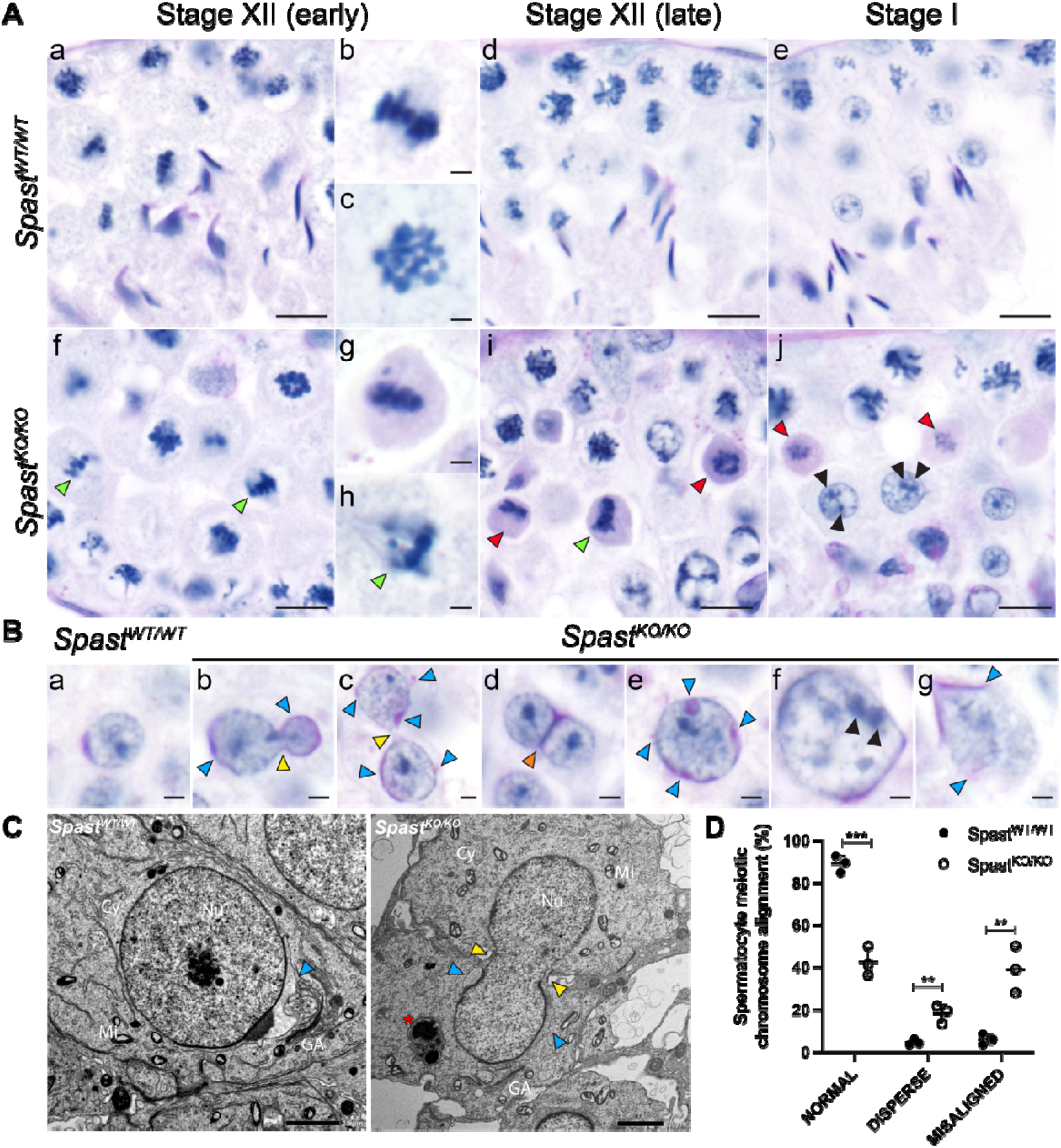
Spastin is essential for correct meiotic division. (**A**) PAS-stained testis sections from *Spast*^*KO/KO*^ mice had an increase in pyknotic spermatocytes (red arrows) in stage XII and I tubules. The green arrowheads indicate wide dispersion or misalignment of chromosomes. The black arrowheads indicate multiple nucleoli within abnormally large nuclei. Panels **b-c** and **g-h** show meiotic cells at increased magnification. Scale bars in **a, d-f, i-j** = 10 µm. Scale bars in **b-c, g-h** = 2 µm. (**B**) The meiosis abnormalities resulted in round spermatids with abnormal phenotypes, including sister cells sharing a single nucleus, which crossed the intercellular bridge (yellow arrowheads), binucleated spermatids (orange arrowhead), and abnormally large nuclei (e-g). Multiple, or fragmented, acrosomes are also indicated by blue arrowheads. Scale bars in **a-g** = 2 µm. (**C**) Failure of meiosis resulting in a nucleus crossing the intercellular bridge between two daughter cells. The position of the intercellular bridge is indicated by yellow arrowheads; the position of the acrosome is indicated by blue arrowheads. The asterisk indicates what is likely to be a late-stage multilamellar body, which may have formed due to an overactive Golgi-apparatus or due to the over-activation of phagocytic pathways. This structure was frequently observed in *Spast*^*KO/KO*^ mice but was not observed in *Spast*^*WT/WT*^ mice. Cytoplasm, Cy; Golgi apparatus, GA; Mitochondrion, Mi; Nucleus, Nu. Scale bars = 2 µm. (**D**) Quantification of common phenotypic defects seen in meiosis in *Spast*^*KO/KO*^ mice (black circles) compared to *Spast*^*WT/WT*^ (white circles) controls (n=3/genotype, lines represent mean ± SD). Asterisks denote different levels of significance; ** p< 0.01, *** p< 0.001.

### Spastin is essential for meiotic spindle formation and function in male germ cells

Consistent with the significant reduction in sperm output, there was a significant increase in apoptotic germ cells in *Spast*^*KO/KO*^ mice compared to *Spast*^*WT/WT*^ littermates (Fig. S1A). The increase in apoptosis occurred primarily in metaphase and early anaphase spermatocytes, suggesting an essential role for spastin in male mammalian meiosis (Fig. S1B,C). Indeed, detailed analysis of *Spast*^*KO/KO*^ PAS-stained testis sections revealed metaphase spermatocytes frequently contained misaligned chromosomes (Fig. 3A, green arrowheads), and/or a wider dispersion of chromosomes at the metaphase plate (Fig. 3A, green arrowhead). These phenotypes were rarely observed in *Spast*^*WT/WT*^ controls. On average, 39% of meiotic cells in *Spast*^*KO/KO*^ mice contained misaligned chromosomes and 18% were abnormally dispersed. In contrast, on average, 6% of meiotic cells in *Spast*^*WT/WT*^ mice were misaligned and 5% were abnormally dispersed (Fig. 3D). In the *Spast*^*KO/KO*^ germ cells that progressed to anaphase uneven chromosome segregation was common (Fig. 3B, panel B).

In *Spast*^*WT/WT*^ testis sections, meiosis I completed in stage XII tubules as expected, and meiosis I or II spermatocytes were not seen in subsequent stage I tubules (Fig. 3A). In contrast, in the *Spast*^*KO/KO*^ testis, many pyknotic PAS-positive/caspase-positive metaphase I and early anaphase I spermatocytes were observed to arrest development in stage XII and persisted in stage I tubules (Fig. 3A, red arrowheads, Fig. S1B,C) indicative of a meiosis arrest followed by germ cell loss. Of the *Spast*^*KO/KO*^ germ cells that completed meiosis, many of the resultant round spermatids were abnormal. In *Spast*^*KO/KO*^ males, round spermatids often had abnormally large nuclei (Fig. 3B, panels e-g) containing multiple nucleoli, or the presence of multiple nuclei within the same cell. In contrast, round spermatids from *Spast*^*WT/WT*^ males (Fig. 3B, panel a) had uniformly sized nuclei with a single nucleolus. The absence of a corresponding number of abnormally small diameter spermatids suggests a major failure of chromosome segregation involving the collapse of at least two sets of chromosomes into a single spermatid nucleus (Fig. 3B). An additional unusual phenotype we frequently observed in *Spast*^*KO/KO*^ mice was a single nucleus crossing the intercellular bridge between sister round spermatids (Fig. 3B, yellow arrowheads). This phenotype was never seen in wild-type mice and is suggestive of increased malleability of mutant cells and a failure of metaphase/anaphase and incomplete cytokinesis. In addition, rare binucleated spermatids were observed in the *Spast*^*KO/KO*^ mice (Fig. 3B, orange arrowhead). These cells likely arose as a result of complete anaphase followed by unsuccessful cytokinesis. Neither of these phenotypes were observed in the round spermatids from *Spast*^*WT/WT*^ mice (Fig. 3B, panel a).

### Spastin is required for acrosome development

Despite the meiotic disruptions observed in *Spast*^*KO/KO*^ testes, the processes that govern the morphogenesis of round spermatids into elongated spermatids continued in *Spast*^*KO/KO*^ germ cells. One of the earliest morphological events is acrosome formation which occurs at the apical surface of the sperm nucleus. This structure is required for the penetration of the cells surrounding the oocyte and thus fertilisation. It begins with the production of pro-acrosomal vesicles in step 2-3 spermatids. In wild-type spermatids these vesicles are transported to the apical pole of the nucleus where they adhere to the nuclear envelope via the acroplaxome to form a single acrosomal vesicle (Fig. 4B). During the Golgi phase (step 2-3) of acrosome development, pro-acrosomal vesicles are solely derived from the Golgi, whereas in the cap phase of development (step 4-7) both Golgi and endocytic pathway-derived vesicles progressively enlarge the acrosome as it flattens and spreads to cover the apical half of the nucleus (Fig. 4B, cap phase) (Pleuger et al., 2020).

**Fig. 4:**
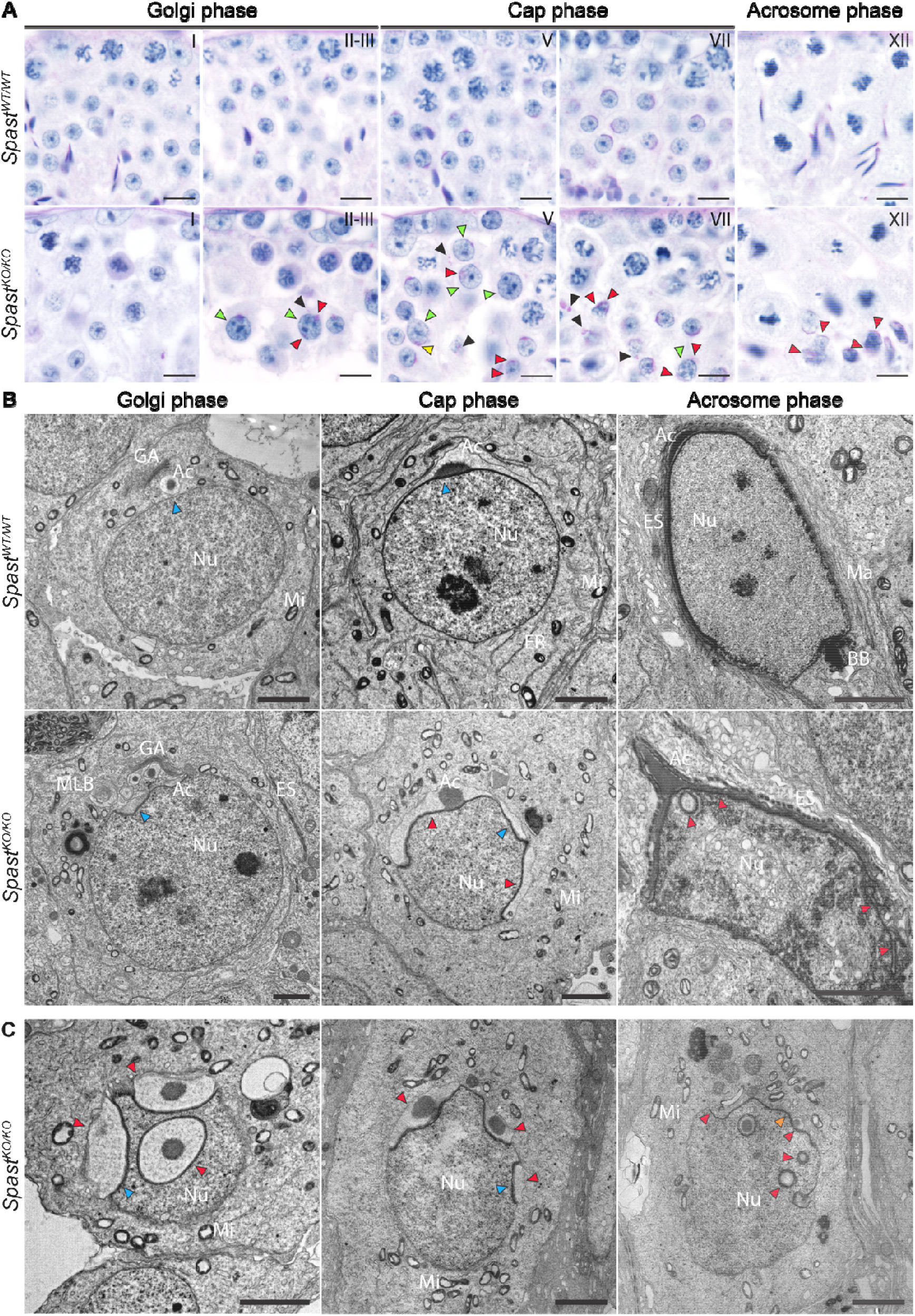
Spastin is essential for the formation of the acrosome. (**A**) The absence of spastin resulted in multiple defects during acrosome development as observed in PAS-stained testis sections. Red arrowheads indicate the presence of multiple pro-acrosomal vesicles, with some having an abnormal localisation within the cytoplasm and others being incorrectly localised at the nucleus. The yellow arrowhead indicates an acrosomal vesicle developing on a nucleus traversing the intercellular bridge, and green arrowheads indicate overtly abnormally large spermatid nuclei. Roman numerals indicate seminiferous tubule stage. Scale bars = 10 µm. (**B**) and (**C**) Transmission electron microscopy showing the ultrastructure of the acrosome in spermatids from *Spast*^*WT/WT*^ and *Spast*^*KO/KO*^ males. In (**B**) progressive steps of acrosome development are shown from left to right. The blue arrowheads indicate an abnormal invagination of the nuclear membrane below the developing acrosomal granules, and lack of this invagination in the *Spast*^*WT/WT*^ mice. Red arrowheads indicate sites of supernumerary acrosome formation. Orange arrowhead indicates abnormal nuclear membrane morphology in the absence of the acrosomal vesicle. Acrosome, Ac; Ectoplasmic specialisation, ES; Golgi Apparatus, GA; Manchette, Ma; Mitochondria, Mi; Multilamellar Body, MLB; Nucleus, Nu. Scale bars = 2 µm.

We observed that early round spermatids from *Spast*^*KO/KO*^ males (step 2-3) had PAS-positive pro-acrosomal vesicles that were ectopically distributed throughout the cytoplasm (Fig. 4A, black arrowheads). In many spermatids, pro-acrosomal vesicles were observed to adhere to multiple ectopic sites on the nuclear membrane, including at the caudal pole (Fig. 4A red arrowheads), suggesting a disruption of the cytoskeletal network required for pro-acrosomal vesicle transport from the Golgi to the nuclear membrane. Moreover, as spermatids from *Spast*^*KO/KO*^ males developed into cap phase, this resulted in supernumerary acrosomes (Fig. 4B-C, red arrowheads). Additionally, multi-lamellar bodies were frequently observed in spermatids from *Spast*^*KO/KO*^ mice from cap phase onwards (Fig. 3C, asterisk) indicating the Golgi apparatus and/or the endocytic pathway may be overactive (Hariri et al., 2000).

In the acrosome phase of development (step 8-12 spermatids), the acrosome could be seen as a thin vesicle coating the entire anterior region of the sperm head in spermatids from *Spast*^*WT/WT*^ males (Fig. 4B, acrosome phase). A similar compacted acrosome phenotype was seen in the spermatids from *Spast*^*KO/KO*^ males, however, multiple acrosome compartments were still observed (Fig. 4B, red arrowheads), in addition to a loss of nuclear membrane integrity (Fig. 4B, green arrowhead). Electron microscopy revealed that docking of acrosomal vesicles throughout development, starting in the Golgi phase, was associated with an abnormally deep nuclear membrane invagination in spermatids from *Spast*^*KO/KO*^ males, suggesting compromised nuclear integrity (Fig. 4B, blue arrowheads).

### Spastin is required for the maintenance of nuclear membrane integrity

One of the more unusual manifestations of spastin loss was a severe disruption to spermatid nuclear integrity at the onset of nuclear elongation in step 9 (Fig. 5E-L). This phenotype was never observed in *Spast*^*WT/WT*^ controls (Fig. 5A-D). Nuclear envelope breakages were first apparent in early elongating spermatids from *Spast*^*KO/KO*^ mice (Fig. 5F, J). At later developmental stages, the nuclear envelopes of spermatids from *Spast*^*KO/KO*^ males became increasingly degraded and the mixing of nuclear and cytoplasmic material continued until they were indistinguishable from each other (Fig. 5E,F,H-L). Ruptured nuclear membranes were most frequently observed at the caudal pole (Fig. 5J, red arrowhead), and were rarely seen at the acrosome-covered apical pole, possibly due to a stabilising effect of the acrosome and/or acroplaxome on the membrane. Alternatively, the pressure applied by the manchette, which envelops the caudal half of the spermatid from step 8/9 onwards may be a trigger for rupture. As described in (Lehti and Sironen, 2016), the manchette is a transient microtubule-based structure that plays a pivotal role in sculpting the distal half of the spermatid nucleus.

**Fig. 5:**
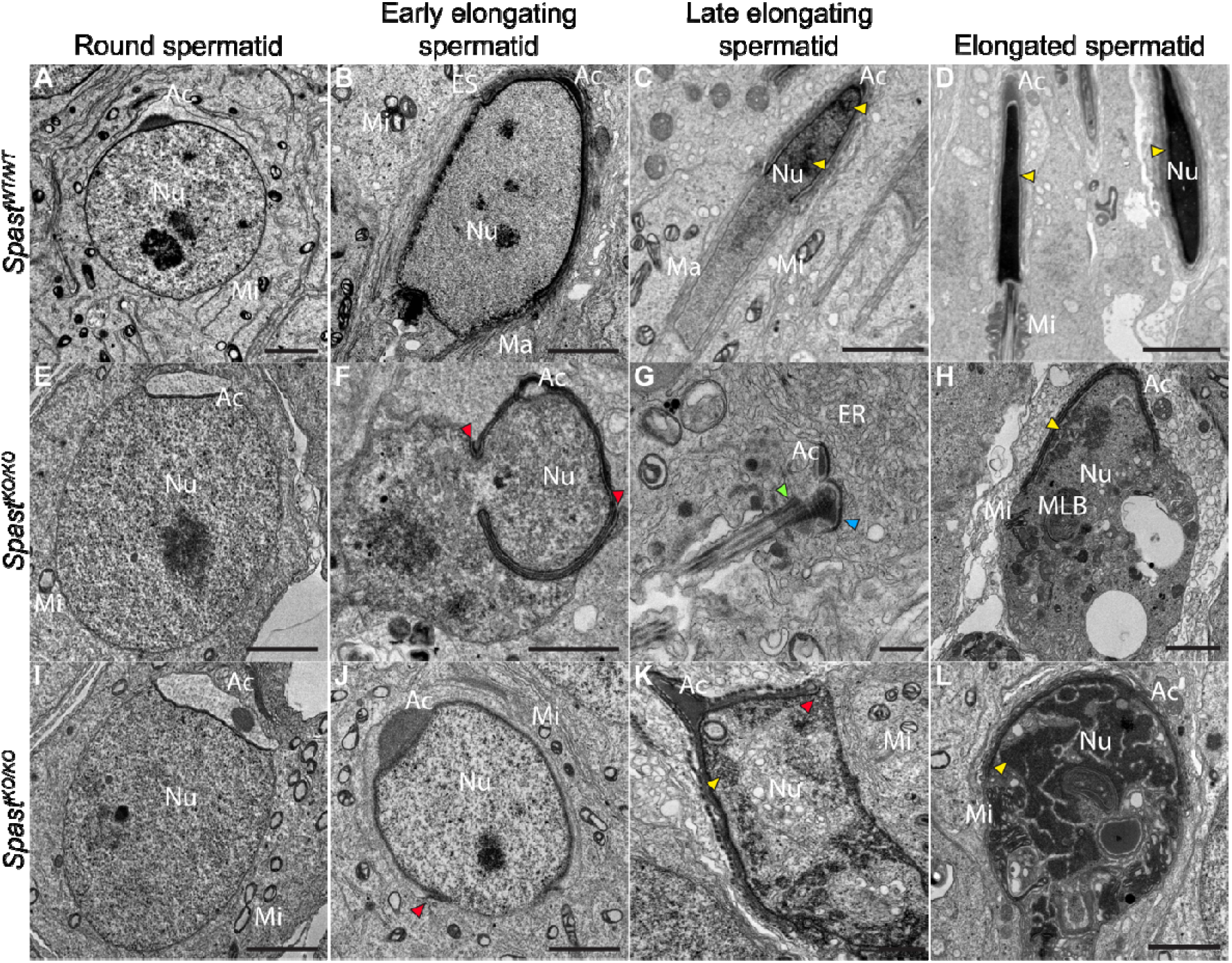
Spastin is required for the maintenance of spermatid nuclear membrane integrity. Transmission electron microscopy of developing spermatids from *Spast*^*WT/WT*^ and *Spast*^*KO/KO*^ mice. In *Spast*^*KO/KO*^ mice, following the initiation of spermatid elongation, spermatids presented with a loss of nuclear membrane integrity ultimately resulting in cell death and a virtual absence of sperm. Red arrowheads indicate the site of membrane rupture. Blue arrowhead indicates the basal plate of the head-tail coupling apparatus. Yellow arrowheads indicate condensed DNA. The green arrowhead indicates the basal body, and the blue arrowhead indicates the associated nuclear membrane. Acrosome, Ac; Endoplasmic reticulum, ER; Ectoplasmic specialisation, ES; Manchette, Ma; Mitochondria, Mi; Multilamellar Body, MLB; Nucleus, Nu. Scale bars **A-F, H**-**L** = 2 µm. Scale bar **G** = 500 nm.

We also noted that DNA condensation was disrupted in spermatids from *Spast*^*KO/KO*^ mice. In elongated spermatids from *Spast*^*WT/WT*^ mice, the nuclear material became progressively more electron dense as DNA condensed (Fig. 5, yellow arrowheads). In elongating spermatids from *Spast*^*KO/KO*^ males, DNA condensation was only initiated in isolated regions (Fig. 5, yellow arrowheads). In the later steps of spermiogenesis, and in contrast to the situation in wild-type, which was replete with elongating and elongated spermatids, these defects collectively resulted in most spermatids from *Spast*^*KO/KO*^ males containing no discernible nucleus (Fig. 5K-L). However, fragments of the nuclear membrane associated with the acrosome (Fig. 5K) and/or the basal body from which the sperm tail initiates (Fig. 5G, blue arrowhead) were visible. Consistent with these defects, spermatids from *Spast*^*KO/KO*^ males had an increase in DNA damage when compared to *Spast* ^*WT/WT*^ as assessed by marking _γ_-H2AX, an indicator of double stranded DNA breaks (Fig. S2)

### Spastin is required for manchette development and sperm head shaping

Spermatid head shaping is mediated in part by the manchette (reviewed in (Dunleavy et al., 2019)), a transient structure made up of microtubules that extend caudally from a perinuclear ring immediately distal to the leading edge of the acrosome. In spermatids from *Spast*^*WT/WT*^ males, the manchette forms at step 8, and as spermatogenesis progresses, the manchette moves distally towards the centriole/basal body and the growing sperm tail (Fig. 6B). In parallel, the perinuclear ring constricts, thus acting to sculpt the distal half of the sperm head (Fig. 6B, Stage XI). Once sperm head shaping is complete, the manchette is disassembled in step 14 spermatids (Fig. 6B, Stage II-III).

**Fig. 6:**
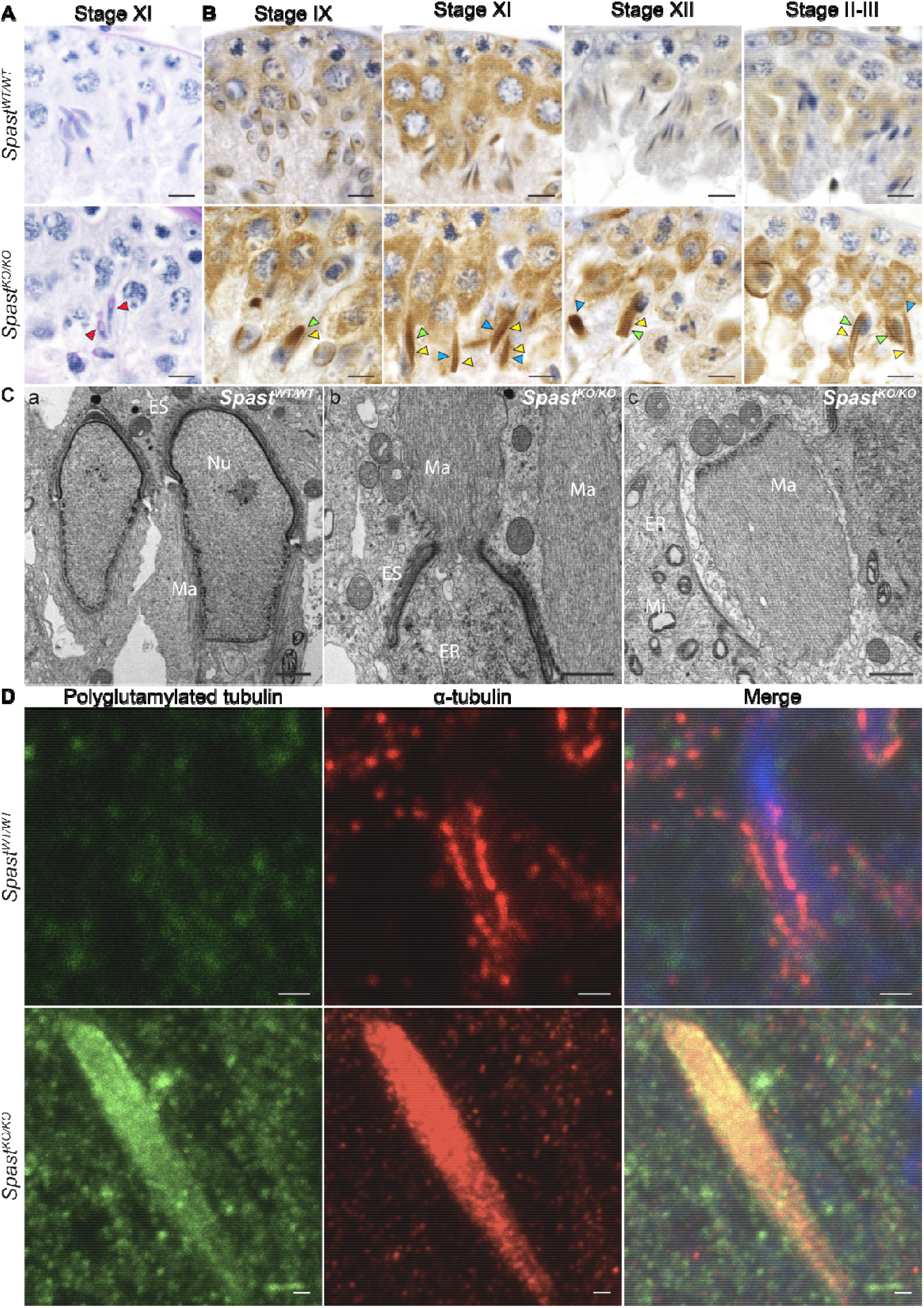
Spastin is a key regulator of manchette structure and dynamics. (**A**) PAS-stained testis sections showing normal elongating spermatids in *Spast*^*WT/WT*^ mice and abnormal elongating spermatids in *Spast*^*KO/KO*^ mouse testes (red arrowheads). (**B**) *Spast*^*WT/WT*^ and *Spast*^*KO/KO*^ testis sections immunolabelled for α-tubulin, a core component of microtubules within the manchette. The tubule stages that capture manchette formation, migration, and disassembly are shown from left to right. Green and blue arrowheads respectively, indicate manchettes that have partially or completely dissociated from the nucleus. The blue arrowheads indicate manchettes of abnormal size. Scale bars of **A-B** = 20 µm. (**C**) TEM showing the manchette ultrastructure in *Spast*^*WT/WT*^ and *Spast*^*KO/KO*^ mice. In panel b, a manchette dissociating from the nucleus of a spermatid in a *Spast*^*KO/KO*^ male can be observed, and in panel c a dissociated manchette is shown. Endoplasmic reticulum, ER; Ectoplasmic specialisation, ES; Manchette, Ma; Mitochondria, Mi; Nucleus, Nu. Scale bars of **C** = 1 µm. (**D**) Immunostaining of manchettes showing an increase in microtubule number (red) and polyglutamylated tubulin (green) in *Spast*^*KO/KO*^ compared to *Spast*^*WT/WT*^ mice. Nuclei are counterstained with DAPI (blue). Staining for polyglutamylated tubulin identified an overall increase in polyglutamylated tubulin in *Spast*^*KO/KO*^ spermatids, especially in the microtubules of the manchette, consistent with polyglutamylated tubulin being the preferred target for spastin-mediated microtubule severing. Scale bars of **D** = 1µm.

In *Spast*^*KO/KO*^ mice, the manchette (marked by _α_-tubulin) formed at the appropriate time, but was phenotypically abnormal (Fig. 6B). Manchette microtubules were observed to over-accumulate resulting in abnormally wide and dense manchettes suggestive of a role for spastin in microtubule pruning (Fig. 6B, yellow arrowheads). Consistent with this interpretation, by step 11, manchettes in *Spast*^*KO/KO*^ males were excessively long compared to those seen in *Spast*^*WT/WT*^ males (Fig. 6B). Further, manchettes were still present in step 14 spermatids from *Spast*^*KO/KO*^ (stage II-III), while in spermatids from *Spast*^*WT/WT*^ males they were disassembled (Fig. 6A, stage II-III), indicating that like katanin proteins (Dunleavy et al., 2021, Dunleavy et al., 2017), spastin influences the dissolution of the manchette. The absence of spastin resulted in the partial (Fig. 6B green arrowheads) or complete detachment (Fig. 6B, blue arrowheads) of the manchette from most elongating spermatid nuclei. This dissociation could be due to the degradation of the nuclear membrane resulting from the compromised nuclear integrity explored above.

Previous studies have shown that spastin preferentially severs polyglutamylated tubulin and that knockout of spastin resulted in an increase of polyglutamylated microtubules (Magiera et al., 2018, Lacroix et al., 2010). As such, we hypothesised that if spastin were required to sever manchette microtubules, there would be an increase of polyglutamylated microtubules in the manchettes of *Spast*^*KO/KO*^ testis sections. To test this, we marked testis sections for α-tubulin and polyglutamylated tubulin. In *Spast*^*KO/KO*^ mice, we found not only was manchette microtubule density increased compared to *Spast*^*WT/WT*^, but there was an increased accumulation of polyglutamylated microtubules (Fig. 6D). Collectively, these results strongly suggest that within male germ cells, spastin severs polyglutamylated microtubules within the manchette to control the number of microtubules within the manchette and to disassemble the manchette at the end of spermiogenesis.

## Discussion

Previously, we have shown that spermatogenesis is critically dependent on katanin-mediated microtubule severing (Dunleavy et al., 2021, Dunleavy et al., 2017, O’Donnell et al., 2012, Smith et al., 2012). Here, we reveal the microtubule severing protein, spastin, is also essential for multiple aspects of male germ development, and that its loss is ultimately incompatible with sperm production (azoospermia), due to meiotic failure followed by a catastrophic loss of nuclear structure. Our data reveals that spastin is an essential regulator of meiosis, wherein it regulates metaphase and anaphase spindle function, and cytokinesis. We also reveal spastin has essential roles in acrosome assembly, in ensuring spermatid nuclear integrity, and in defining the structure and function of the manchette.

While no defects were apparent during *Spast*^*KO/KO*^ male germ cell mitosis, our data establishes spastin as essential for the correct completion of male meiosis in mice. In the absence of spastin, we found increases in chromosome misalignment at the metaphase plate, in failed chromosome segregation during anaphase, and in failed or abnormal cytokinesis. Chromosome misalignment at metaphase could be due to a defect in the regulation of the length of the microtubules that make up the bipolar spindle.

During anaphase, the failure of poleward chromosome segregation in *Spast*^*KO/KO*^ spermatocytes, suggests spastin-mediated microtubule severing is required for the poleward shortening of spindle microtubules. This is consistent with in vitro data from mitosis in *D. melanogaster*, wherein spastin was shown to promote poleward chromosome movement during anaphase ‘Pacman flux’, by stimulating depolymerisation of microtubule minus-ends at the spindle pole (Zhang et al., 2007). Of note, this work also identified a parallel role for the microtubule-severing enzyme fidgetin in severing at the spindle poles during ‘Pacman flux’ (Zhang et al., 2007). It is thus possible that that in mammals, the fidgetins can compensate for spastin function during mitosis anaphase, but not in meiosis. Spastin has previously been found to localise to the spindle poles in HeLa cells, supporting a role of spastin mediated severing in regulating the bipolar spindle in mammals (Errico et al., 2004).

The occurrences of binucleated spermatids in the absence of spastin, suggest that spastin-mediated microtubule severing is required to regulate midbody microtubule abscission during meiosis. Indeed, this is consistent with data showing spastin severs midbody microtubules during mitosis in HeLa cells (Connell et al., 2009, Yang et al., 2008, Pisciottani et al., 2019). More commonly however, in the absence of spastin we observed a single ‘pinched’ spermatid nucleus shared by two sister cells across an intercellular bridge. The likely explanation for this phenotype is that the ‘pinched’ nucleus occurs when there is a failure of anaphase resulting in a single large round spermatid nucleus, and then cytokinesis proceeds regardless of the position and size of the nucleus. The role spastin-mediated severing plays in establishing nuclear integrity may allow these nuclei to be flexible enough for the pinched phenotype to occur. Interestingly, our previous work identified frequent occurrences of binucleated spermatids when katanin was lost, but never the ‘pinched’ nuclear phenotype (Dunleavy et al., 2021) indicating that spastin is essential at additional stages of anaphase and/or cytokinesis in male meiosis.

In *Spast*^*KO/KO*^ mice, we observed spermatid nuclei that had compromised membrane integrity. Compromised nuclear integrity first presented as deep invaginations of the nuclear membrane caused by the overlying acrosome and progressed to ruptured nuclei after manchette formation. We predict compromised nuclear integrity was due to the requirement of spastin and ESCRT-III components for nuclear membrane fission following cell division, as shown in HeLa cells (Vietri et al., 2015). Specifically, we hypothesise that the progressive loss of nuclear integrity seen in post-meiotic spermatids is due to the inability of the compromised nuclear membrane to withstand the pressure produced by the events of spermiogenesis. Consistent with the results of Vietri et al., we observed an increase in DNA double-stranded breaks in *Spast*^*KO/KO*^ mice after meiosis.

Our data also reveals spastin is required for the correct localisation and assembly of the acrosome during development. Without spastin, pro-acrosomal vesicles were mis-trafficked to the cytoplasm or to ectopic locations on the nuclear envelope. While difficult to see at the light microscope level, this defect may be due to an increase in the number of microtubules emanating from the Golgi apparatus throughout the cytoplasm. Therefore, we propose that spastin is required to prune microtubule tracks not typically used in pro-acrosomal vesicle trafficking. Consistent with this, previous work on mouse neuronal development found that spastin is required for the pruning of axon branches during neurogenesis (Brill et al., 2016), and that microtubule-based axonal transport is disrupted in the absence of spastin (Tarrade et al., 2006).

Finally, we reveal that spastin is an essential regulator of manchette microtubule density, length, and disassembly and manchette-nuclear attachment. Previous research found that spastin preferentially severs polyglutamylated microtubules (Lacroix et al., 2010, Valenstein and Roll-Mecak, 2016) and that loss of spastin leads to an increase in polyglutamylated tubulin (Magiera et al., 2018). Our results support a role for spastin in regulating the accumulation and length of manchette microtubules as we found an increase in polyglutamylated microtubules within *Spast*^*KO/KO*^ manchettes. Our previous work identified that the katanin, KATNAL2 is also important for manchette movement and length, indicating that a suite of microtubule severing enzymes are required to regulate different aspects of the manchette (Dunleavy et al., 2017).

To our knowledge, this is the first time that many *in vitro* phenotypes resulting from a loss of spastin have been confirmed in an *in vivo* model. We have shown a phenotype consistent with work showing that spastin is required for the completion of nuclear envelope reformation and for midbody abscission and have highlighted the biological relevance of these roles during spermatogenesis. Additionally, aspects of the manchette phenotype observed are unique and suggest a distinct role for spastin in the regulation of complex microtubule-based structures. This work has established spastin as a key regulator of microtubule dynamics during spermatogenesis, and many microtubule-dependent processes are disrupted without its action.

Our work provides a better understanding of disrupted cell dynamics in cases of hereditary spastic paraplegia, increases our understanding of the role of microtubule regulation in spermatogenesis and may ultimately inform fertility care for patients carrying SPAST loss-of-function genetic variants.

## Materials and Methods

### Animal ethics statement

All animal procedures were performed with the approval of the Monash University Animal Experimentation Ethics Committee or the University of Melbourne Ethics Committee and were consistent with the requirements set out in the Australian National Health and Medical Research Council (NHMRC) Guidelines on Ethics in Animal Experimentation.

### Mouse model production and phenotypic analysis

The mouse model used for this study was first described in (Brill et al., 2016) where they showed that spastin was involved in dendritic pruning. In brief, the *Spast*^*tm1a(KOMP)Wtsi*^ targeting vector (PG00198_Z_2_G10) was generated by the trans-NIH Knockout Mouse Project (www.komp.org). The construct contains a splice site acceptor and a poly-adenylation sequence resulting in truncation following exon 4 (ENSMUSE00000137944) of *Spast* (ENSMUSG00000024068). Mice were maintained on a C57BL/6 background. Wild-type littermates (*Spast*^*WT/WT*^) were used as controls for *Spast* knockout mice (*Spast*^*KO/KO*^), and all male mice used for analysis were adult (≥ 10 weeks of age). Mouse genotypes were identified from tail biopsies using real-time PCR with probes designed for each allele (Transnetyx, Cordova, TN). *Spast* mRNA levels in *Spast*^*KO/KO*^ mice were tested using qPCR on whole testis tissue as described below. Protein levels were determined by western blotting of testis lysates as described below.

### Quantitative qPCR

Whole testes were homogenized, and total RNA was extracted using Trizol reagent (Life Technologies) and cDNA synthesised using SuperScript III reverse transcriptase (Life Technologies). To verify the truncation of the *Spast* gene in the *Spast*^*KO/KO*^ mouse line, PCR primer sets were designed that span part of exon 4 and exon 5 (forward 5’-TAACCTGACATGCCGCAATG-3’ and reverse 5’-ACAAACCACTGCAACTAGGC-3’). qPCR was performed using SYBR Select Master Mix (Applied Biosystems). Each reaction was performed in triplicate, and on three biological replicates per genotype, on an Applied Biosystems QuantStudio 3 real-time PCR system. DNA was denatured at 95°C for two minutes, followed by 35 cycles of 95°C for 30 seconds and then 60°C for one minute. *Ppia* was amplified simultaneously as an internal control (forward primer 5’-CGTCTCCTTCGAGCTGTTT-’3 and reverse primer 5’-CCCTGGCACATGAATCCT-’3). All results were normalised to the expression of *Ppia*. Differential expression was analysed using the 2ΔΔ^CT^ method (Livak and Schmittgen, 2001).

### Fertility Characterisation

The fertility of the *Spast*^*KO/KO*^ mouse line was characterised as described in (Houston et al., 2021). Fertility tests used male mice ≥ 10 weeks of age, in which males were mated with two wild-type females (≥ 6 weeks of age). Females were monitored for copulatory plugs as an indication of successful mating and the number of pups born per copulatory plug was recorded. Testis daily sperm production (DSP) and total epididymal sperm content (n≥3 mice / genotype) were determined using the Triton X-100 nuclear solubilisation method described in (Dunleavy et al., 2021).

Testes and epididymides were fixed in Bouin’s fixative and processed into paraffin for histological examination. Periodic acid-Schiff (PAS) and haematoxylin staining was used to visualise male reproductive tract histology (n≥3 mice / genotype). Germ cell apoptosis was evaluated by immunostaining for cleaved Caspases 3 and 9 and counterstaining for haematoxylin as previously described (O’Bryan et al., 2013). The number of caspase-positive cells in a minimum of 100 randomly selected seminiferous tubules per mouse was quantified and statistical analysis was performed as detailed below (n = 3 mice / genotype).

### Transmission electron microscopy

To analyse the ultrastructure of the seminiferous tubules, partially decapsulated testes were processed for transmission electron microscopy as in (Dunleavy et al., 2021). Ultrathin sections were cut on a Reichert Jung Ultracut Microtome and placed on 100×100 square copper grids (ProSciTech). Sections were analysed using a Jeol JEM-1400 Plus transmission electron microscope at the Monash University Ramaciotti Centre for Cryo-Electron Microscopy (Monash University, Clayton).

### Antibodies

Primary antibodies used included those against α-tubulin (T5168, Sigma, ascites fluid, 1 in 5000 and ab4074, Abcam, 1 μg ml^-1^) polyglutamylated tubulin B3 (T9822, Sigma, 2 µg ml^-1^), γH2AX (05-636, Millipore, 0.1 µg ml^-1^), cleaved caspase 3 (9664, Cell Signalling, 0.5 μg ml^-1^) and cleaved-caspase 9 (9509, Cell Signaling, 1 μg ml^-1^). Secondary antibodies included Alexa Fluor 488 donkey anti-goat (A11055, Invitrogen), Alexa Fluor 555 donkey anti-goat (A21432, Invitrogen), Alexa Fluor 555 donkey anti-mouse (A31570, Invitrogen), Alexa Fluor 647 donkey anti-mouse (A31571, Invitrogen), Alexa Fluor 647 donkey anti-rabbit (A31573, Invitrogen). Parallel sections were processed in the absence of a primary antibody to control for secondary antibody specificity.

### Immunochemistry

Five micrometre sections were cut from paraffin blocks and dewaxed prior to antigen retrieval by microwaving the sections in 10 mM citrate buffer (pH 6.0) for 16 minutes as previously described ((Jamsai et al., 2008). For colourimetric immunohistochemistry, endogenous peroxidase activity was blocked with 3% H_2_O_2_ in H_2_O for five minutes, and non-specific antibody binding was minimised by blocking with CAS block (Invitrogen) for at least 30 minutes. Primary antibodies were diluted in Dako antibody diluent (S0809, Dako) and incubated overnight at 4°C. Dako envision polymer Dual link system-HRP (K4063, Dako) was applied undiluted for one hour at room temperature. Dako liquid DAB+ substrate chromogen (K3468, Dako) was applied to samples for one minute followed by immediate submersion in water. Sections were counterstained with haematoxylin then dehydrated and mounted with DPX (44581, Sigma-Aldrich).

For immunofluorescence labelling, after dewaxing and antigen retrieval non-specific antibody binding was minimised by incubating sections in CAS Block (Invitrogen). Primary antibodies were diluted in Dako antibody diluent (S0809, Dako) and incubated on sections overnight at 4°C. Secondary antibodies were diluted 1 in 500 in PBS and incubated on sections at room temperature for one hour. DNA was visualized using 1 µg ml^-1^ 4’,6-diamidino-2-phenylindole (DAPI, Invitrogen). Acrosomes were visualized using 0.5 µg ml^-1^ lectin peanut agglutinin (PNA) Alexa Fluor 488 conjugate (L21409, Life Technologies). Sections were mounted under Dako fluorescence mounting medium and glass coverslips (GM304, Dako).

Immunofluorescent images were taken with a Leica TCS SP8 confocal microscope (Leica Microsystems) at the University of Melbourne Biological Optical Microscopy Platform. All images were taken using the 63x/1.40 HC PL APO CS2 oil immersion objective. Z-stacks of testis sections were collected at 0.3µm intervals and assembled into maximum intensity projections in ImageJ were processed using ImageJ 2.1.0.

### Statistics and Reproducibility

Statistical analysis of the germ cell apoptosis data was performed in R version 3.5.1 (R Core Team, 2014). Generalised linear mixed models (GLM) were used to compare the number of caspase-positive cells per tubule between genotypes. For each model, Akaike information criterion (AIC) estimates were used to select the most appropriate error distribution and link functions (i.e., poisson, negative binomial, zero-inflated poisson, zero-inflated negative binomial) using the glmer function (lme4 package; (Bates et al., 2015)) and the glmmTMB function (glmmTMB package; (Brooks et al., 2017)). For all models, a zero-inflated negative binomial distribution (fitted with glmmTMB, using the ziformula argument) was selected as the most appropriate error distribution and link function (i.e., had the lowest AIC score).

All other statistical analysis was performed in GraphPad Prism version 9.3. The statistical significance of differences between two groups was determined using an unpaired student’s T test, significance was defined as p-value < 0.05. For each group a minimum of n = 3 individuals per group were analysed.

## Acknowledgments

We thank the Monash Histology Platform, the Monash Ramaciotti Centre for Cryo-Electron Microscopy, the Monash Animal Research Platform, and the University of Melbourne Biological Optical Microscopy Platform for technical support.

## References

Ahmad, F. J., Yu, W., Mcnally, F. J. & Baas, P. W. 1999. An essential role for katanin in severing microtubules in the neuron. J Cell Biol, 145, 305–15.

Allison, R., Edgar, J. R., Pearson, G., Rizo, T., Newton, T., Gunther, S., Berner, F., Hague, J., Connell, J. W., Winkler, J., Lippincott-Schwartz, J., Beetz, C., Winner, B. & Reid, E. 2017. Defects in ER-endosome contacts impact lysosome function in hereditary spastic paraplegia. J Cell Biol, 216, 1337–1355.

Allison, R., Edgar, J. R. & Reid, E. 2019. Spastin MIT Domain Disease-Associated Mutations Disrupt Lysosomal Function. Front Neurosci, 13, 1179.

Allison, R., Lumb, J. H., Fassier, C., Connell, J. W., Ten Martin, D., Seaman, M. N., Hazan, J. & Reid, E. 2013. An ESCRT-spastin interaction promotes fission of recycling tubules from the endosome. J Cell Biol, 202, 527–43.

Arribat, Y., Grepper, D., Lagarrigue, S., Qi, T., Cohen, S. & Amati, F. 2020. Spastin mutations impair coordination between lipid droplet dispersion and reticulum. PLoS Genet, 16, e1008665.

Bates, D., Machler, M., Bolker, B. M. & Walker, S. C. 2015. Fitting Linear Mixed-Effects Models Using lme4. Journal of Statistical Software, 67, 1–48.

Brill, M. S., Kleele, T., Ruschkies, L., Wang, M., Marahori, N. A., Reuter, M. S., Hausrat, T. J., Weigand, E., Fisher, M., Ahles, A., Engelhardt, S., Bishop, D. L., Kneussel, M. & Misgeld, T. 2016. Branch-Specific Microtubule Destabilization Mediates Axon Branch Loss during Neuromuscular Synapse Elimination. Neuron, 92, 845–856.

Brooks, M. E., Kristensen, K., Van Benthem, K. J., Magnusson, A., Berg, C. W., Nielsen, A., Skaug, H. J., Machler, M. & Bolker, B. M. 2017. glmmTMB Balances Speed and Flexibility Among Packages for Zero-inflated Generalized Linear Mixed Modeling. R Journal, 9, 378–400.

Chang, C. L., Weigel, A. V., Ioannou, M. S., Pasolli, H. A., Xu, C. S., Peale, D. R., Shtengel, G., Freeman, M., Hess, H. F., Blackstone, C. & Lippincott-Schwartz, J. 2019. Spastin tethers lipid droplets to peroxisomes and directs fatty acid trafficking through ESCRT-III. J Cell Biol, 218, 2583–2599.

Chen, K., Ye, Y., Ji, Z., Tan, M., Li, S., Zhang, J., Guo, G. & Lin, H. 2014. Katanin p60 promotes neurite growth and collateral formation in the hippocampus. Int J Clin Exp Med, 7, 2463–70.

Claudiani, P., Riano, E., Errico, A., Andolfi, G. & Rugarli, E. I. 2005. Spastin subcellular localization is regulated through usage of different translation start sites and active export from the nucleus. Experimental Cell Research, 309, 358–369.

Connell, J. W., Allison, R. & Reid, E. 2016. Quantitative Gait Analysis Using a Motorized Treadmill System Sensitively Detects Motor Abnormalities in Mice Expressing ATPase Defective Spastin. PLoS One, 11, e0152413.

Connell, J. W., Lindon, C., Luzio, J. P. & Reid, E. 2009. Spastin couples microtubule severing to membrane traffic in completion of cytokinesis and secretion. Traffic, 10, 42–56.

Dunleavy, J. E. M., O’Bryan, M. K., Stanton, P. G. & O’Donnell, L. 2019. The cytoskeleton in spermatogenesis. Reproduction, 157, R53–R72.

Dunleavy, J. E. M., O’Connor, A. E., Okuda, H., Merriner, D. J. & O’Bryan, M. K. 2021. KATNB1 is a master regulator of multiple katanin enzymes in male meiosis and haploid germ cell development. Development, 148.

Dunleavy, J. E. M., Okuda, H., O’Connor, A. E., Merriner, D. J., O’Donnell, L., Jamsai, D., Bergmann, M. & O’Bryan, M. K. 2017. Katanin-like 2 (KATNAL2) functions in multiple aspects of haploid male germ cell development in the mouse. PLoS Genet, 13, e1007078.

Erdmann, R., Wiebel, F. F., Flessau, A., Rytka, J., Beyer, A., Frohlich, K. U. & Kunau, W. H. 1991. PAS1, a yeast gene required for peroxisome biogenesis, encodes a member of a novel family of putative ATPases. Cell, 64, 499–510.

Errico, A., Ballabio, A. & Rugarli, E. I. 2002. Spastin, the protein mutated in autosomal dominant hereditary spastic paraplegia, is involved in microtubule dynamics. Hum Mol Genet, 11, 153–63.

Errico, A., Claudiani, P., D’Addio, M. & Rugarli, E. I. 2004. Spastin interacts with the centrosomal protein NA14, and is enriched in the spindle pole, the midbody and the distal axon. Hum Mol Genet, 13, 2121–32.

Frickey, T. & Lupas, A. N. 2004. Phylogenetic analysis of AAA proteins. J Struct Biol, 146, 2–10.

Hariri, M., Millane, G., Guimond, M. P., Guay, G., Dennis, J. W. & Nabi, I. R. 2000. Biogenesis of multilamellar bodies via autophagy. Mol Biol Cell, 11, 255–68.

Hazan, J., Fonknechten, N., Mavel, D., Paternotte, C., Samson, D., Artiguenave, F., Davoine, C. S., Cruaud, C., Durr, A., Wincker, P., Brottier, P., Cattolico, L., Barbe, V., Burgunder, J. M., Prud’Homme, J. F., Brice, A., Fontaine, B., Heilig, R. & Weissenbach, J. 1999. Spastin, a new AAA protein, is altered in the most frequent form of autosomal dominant spastic paraplegia. Nature Genetics, 23, 296–303.

Houston, B. J., Conrad, D. F. & O’Bryan, M. K. 2021. A framework for high-resolution phenotyping of candidate male infertility mutants: from human to mouse. Hum Genet, 140, 155–182.

Hu, W. F., Pomp, O., Ben-Omran, T., Kodani, A., Henke, K., Mochida, G. H., Yu, T. W., Woodworth, M. B., Bonnard, C., Raj, G. S., Tan, T. T., Hamamy, H., Masri, A., Shboul, M., Al Saffar, M., Partlow, J. N., Al-Dosari, M., Alazami, A., Alowain, M., Alkuraya, F. S., Reiter, J. F., Harris, M. P., Reversade, B. & Walsh, C. A. 2014. Katanin p80 regulates human cortical development by limiting centriole and cilia number. Neuron, 84, 1240–57.

Jamsai, D., Bianco, D. M., Smith, S. J., Merriner, D. J., Ly-Huynh, J. D., Herlihy, A., Niranjan, B., Gibbs, G. M. & O’Bryan, M. K. 2008. Characterization of gametogenetin 1 (GGN1) and its potential role in male fertility through the interaction with the ion channel regulator, cysteine-rich secretory protein 2 (CRISP2) in the sperm tail. Reproduction, 135, 751–9.

Jeong, B., Kim, T. H., Kim, D. S., Shin, W. H., Lee, J. R., Kim, N. S. & Lee, D. Y. 2019. Spastin Contributes to Neural Development through the Regulation of Microtubule Dynamics in the Primary Cilia of Neural Stem Cells. Neuroscience, 411, 76–85.

Ji, Z., Zhang, G., Chen, L., Li, J., Yang, Y., Cha, C., Zhang, J., Lin, H. & Guo, G. 2018. Spastin Interacts with CRMP5 to Promote Neurite Outgrowth by Controlling the Microtubule Dynamics. Dev Neurobiol, 78, 1191–1205.

Karlsson, M., Zhang, C., Mear, L., Zhong, W., Digre, A., Katona, B., Sjostedt, E., Butler, L., Odeberg, J., Dusart, P., Edfors, F., Oksvold, P., Von Feilitzen, K., Zwahlen, M., Arif, M., Altay, O., Li, X., Ozcan, M., Mardinoglu, A., Fagerberg, L., Mulder, J., Luo, Y., Ponten, F., Uhlen, M. & Lindskog, C. 2021. A single-cell type transcriptomics map of human tissues. Sci Adv, 7, eabh2169.

Kasher, P. R., De Vos, K. J., Wharton, S. B., Manser, C., Bennett, E. J., Bingley, M., Wood, J. D., Milner, R., Mcdermott, C. J., Miller, C. C., Shaw, P. J. & Grierson, A. J. 2009. Direct evidence for axonal transport defects in a novel mouse model of mutant spastin-induced hereditary spastic paraplegia (HSP) and human HSP patients. J Neurochem, 110, 34–44.

Lacroix, B., Van Dijk, J., Gold, N. D., Guizetti, J., Aldrian-Herrada, G., Rogowski, K., Gerlich, D. W. & Janke, C. 2010. Tubulin polyglutamylation stimulates spastin-mediated microtubule severing. J Cell Biol, 189, 945–54.

Lehti, M. S. & Sironen, A. 2016. Formation and function of the manchette and flagellum during spermatogenesis. Reproduction, 151, R43–54.

Leo, L., Weissmann, C., Burns, M., Kang, M., Song, Y., Qiang, L., Brady, S. T., Baas, P. W. & Morfini, G. 2017. Mutant spastin proteins promote deficits in axonal transport through an isoform-specific mechanism involving casein kinase 2 activation. Hum Mol Genet, 26, 2321–2334.

Liu, Q., Zhang, G., Ji, Z. & Lin, H. 2021. Molecular and cellular mechanisms of spastin in neural development and disease (Review). Int J Mol Med, 48, 218.

Livak, K. J. & Schmittgen, T. D. 2001. Analysis of relative gene expression data using real-time quantitative PCR and the 2(-Delta Delta C(T)) Method. Methods, 25, 402–8.

Magiera, M. M., Bodakuntla, S., Ziak, J., Lacomme, S., Marques Sousa, P., Leboucher, S., Hausrat, T. J., Bosc, C., Andrieux, A., Kneussel, M., Landry, M., Calas, A., Balastik, M. & Janke, C. 2018. Excessive tubulin polyglutamylation causes neurodegeneration and perturbs neuronal transport. EMBO J, 37.

Mancuso, G. & Rugarli, E. I. 2008. A cryptic promoter in the first exon of the SPG4 gene directs the synthesis of the 60-kDa spastin isoform. BMC Biol, 6, 31.

Mcnally, F. J. & Roll-Mecak, A. 2018. Microtubule-severing enzymes: From cellular functions to molecular mechanism. J Cell Biol, 217, 4057–4069.

Migliano, S. M., Wenzel, E. M. & Stenmark, H. 2022. Biophysical and molecular mechanisms of ESCRT functions, and their implications for disease. Current Opinion in Cell Biology, 75, 102062.

O’Donnell, L., Rhodes, D., Smith, S. J., Merriner, D. J., Clark, B. J., Borg, C., Whittle, B., O’Connor, A. E., Smith, L. B., Mcnally, F. J., De Kretser, D. M., Goodnow, C. C., Ormandy, C. J., Jamsai, D. & O’Bryan, M. K. 2012. An essential role for katanin p80 and microtubule severing in male gamete production. PLoS Genet, 8, e1002698.

O’Bryan, M. K., Clark, B. J., Mclaughlin, E. A., D’Sylva, R., O’Donnell, L., Wilce, J. A., Sutherland, J. M., O’Connor, A. E., Whittle, B., Goodnow, C. C., Ormandy, C. J. & Jamsai, D. 2013. RBM5 Is a Male Germ Cell Splicing Factor and Is Required for Spermatid Differentiation and Male Fertility. PLoS Genetics, 9.

Park, S. H., Zhu, P. P., Parker, R. L. & Blackstone, C. 2010. Hereditary spastic paraplegia proteins REEP1, spastin, and atlastin-1 coordinate microtubule interactions with the tubular ER network. J Clin Invest, 120, 1097–110.

Pisciottani, A., Biancolillo, L., Ferrara, M., Valente, D., Sardina, F., Monteonofrio, L., Camerini, S., Crescenzi, M., Soddu, S. & Rinaldo, C. 2019. HIPK2 Phosphorylates the Microtubule-Severing Enzyme Spastin at S268 for Abscission. Cells, 8.

Pleuger, C., Lehti, M. S., Dunleavy, J. E., Fietz, D. & O’Bryan, M. K. 2020. Haploid male germ cells-the Grand Central Station of protein transport. Hum Reprod Update, 26, 474–500.

Qiang, L., Piermarini, E., Muralidharan, H., Yu, W., Leo, L., Hennessy, L. E., Fernandes, S., Connors, T., Yates, P. L., Swift, M., Zholudeva, L. V., Lane, M. A., Morfini, G., Alexander, G. M., Heiman-Patterson, T. D. & Baas, P. W. 2019. Hereditary spastic paraplegia: gain-of-function mechanisms revealed by new transgenic mouse. Hum Mol Genet, 28, 1136–1152.

Reid, E., Connell, J., Edwards, T. L., Duley, S., Brown, S. E. & Sanderson, C. M. 2005. The hereditary spastic paraplegia protein spastin interacts with the ESCRT-III complex-associated endosomal protein CHMP1B. Hum Mol Genet, 14, 19–38.

Rigden, D. J., Liu, H., Hayes, S. D., Urbe, S. & Clague, M. J. 2009. Ab initio protein modelling reveals novel human MIT domains. FEBS Lett, 583, 872–8.

Roll-Mecak, A. & Vale, R. D. 2008. Structural basis of microtubule severing by the hereditary spastic paraplegia protein spastin. Nature, 451, 363–7.

Sandate, C. R., Szyk, A., Zehr, E. A., Lander, G. C. & Roll-Mecak, A. 2019. An allosteric network in spastin couples multiple activities required for microtubule severing. Nat Struct Mol Biol, 26, 671–678.

Smith, L. B., Milne, L., Nelson, N., Eddie, S., Brown, P., Atanassova, N., O’Bryan, M. K., O’Donnell, L., Rhodes, D., Wells, S., Napper, D., Nolan, P., Lalanne, Z., Cheeseman, M. & Peters, J. 2012. KATNAL1 Regulation of Sertoli Cell Microtubule Dynamics Is Essential for Spermiogenesis and Male Fertility. Plos Genetics, 8.

Snider, J., Thibault, G. & Houry, W. A. 2008. The AAA+ superfamily of functionally diverse proteins. Genome Biol, 9, 216.

Tan, D., Zhang, H., Deng, J., Liu, J., Wen, J., Li, L., Wang, X., Pan, M., Hu, X. & Guo, J. 2020. RhoA-GTPase Modulates Neurite Outgrowth by Regulating the Expression of Spastin and p60-Katanin. Cells, 9.

Tarrade, A., Fassier, C., Courageot, S., Charvin, D., Vitte, J., Peris, L., Thorel, A., Mouisel, E., Fonknechten, N., Roblot, N., Seilhean, D., Dierich, A., Hauw, J. J. & Melki, J. 2006. A mutation of spastin is responsible for swellings and impairment of transport in a region of axon characterized by changes in microtubule composition. Hum Mol Genet, 15, 3544–58.

Vajjhala, P. R., Nguyen, C. H., Landsberg, M. J., Kistler, C., Gan, A. L., King, G. F., Hankamer, B. & Munn, A. L. 2008. The Vps4 C-terminal helix is a critical determinant for assembly and ATPase activity and has elements conserved in other members of the meiotic clade of AAA ATPases. FEBS J, 275, 1427–1449.

Valenstein, M. L. & Roll-Mecak, A. 2016. Graded Control of Microtubule Severing by Tubulin Glutamylation. Cell, 164, 911–21.

Vemu, A., Szczesna, E., Zehr, E. A., Spector, J. O., Grigorieff, N., Deaconescu, A. M. & Roll-Mecak, A. 2018. Severing enzymes amplify microtubule arrays through lattice GTP-tubulin incorporation. Science, 361.

Vietri, M., Schink, K. O., Campsteijn, C., Wegner, C. S., Schultz, S. W., Christ, L., Thoresen, S. B., Brech, A., Raiborg, C. & Stenmark, H. 2015. Spastin and ESCRT-III coordinate mitotic spindle disassembly and nuclear envelope sealing. Nature, 522, 231–5.

White, S. R., Evans, K. J., Lary, J., Cole, J. L. & Lauring, B. 2007. Recognition of C-terminal amino acids in tubulin by pore loops in Spastin is important for microtubule severing. J Cell Biol, 176, 995–1005.

Wood, J. D., Landers, J. A., Bingley, M., Mcdermott, C. J., Thomas-Mcarthur, V., Gleadall, L. J., Shaw, P. J. & Cunliffe, V. T. 2006. The microtubule-severing protein Spastin is essential for axon outgrowth in the zebrafish embryo. Hum Mol Genet, 15, 2763–71.

Yang, D., Rismanchi, N., Renvoise, B., Lippincott-Schwartz, J., Blackstone, C. & Hurley, J. H. 2008. Structural basis for midbody targeting of spastin by the ESCRT-III protein CHMP1B. Nat Struct Mol Biol, 15, 1278–86.

Yu, W., Qiang, L., Solowska, J. M., Karabay, A., Korulu, S. & Baas, P. W. 2008. The microtubule-severing proteins spastin and katanin participate differently in the formation of axonal branches. Mol Biol Cell, 19, 1485–98.

Zehr, E. A., Szyk, A., Szczesna, E. & Roll-Mecak, A. 2020. Katanin Grips the beta-Tubulin Tail through an Electropositive Double Spiral to Sever Microtubules. Dev Cell, 52, 118–131 e6.

Zhang, D., Rogers, G. C., Buster, D. W. & Sharp, D. J. 2007. Three microtubule severing enzymes contribute to the “Pacman-flux” machinery that moves chromosomes. Journal of Cell Biology, 177, 231–242.

