## Supplemental File for "Spastin is an essential regulator of male meiosis, acrosome formation, manchette structure and nuclear integrity"

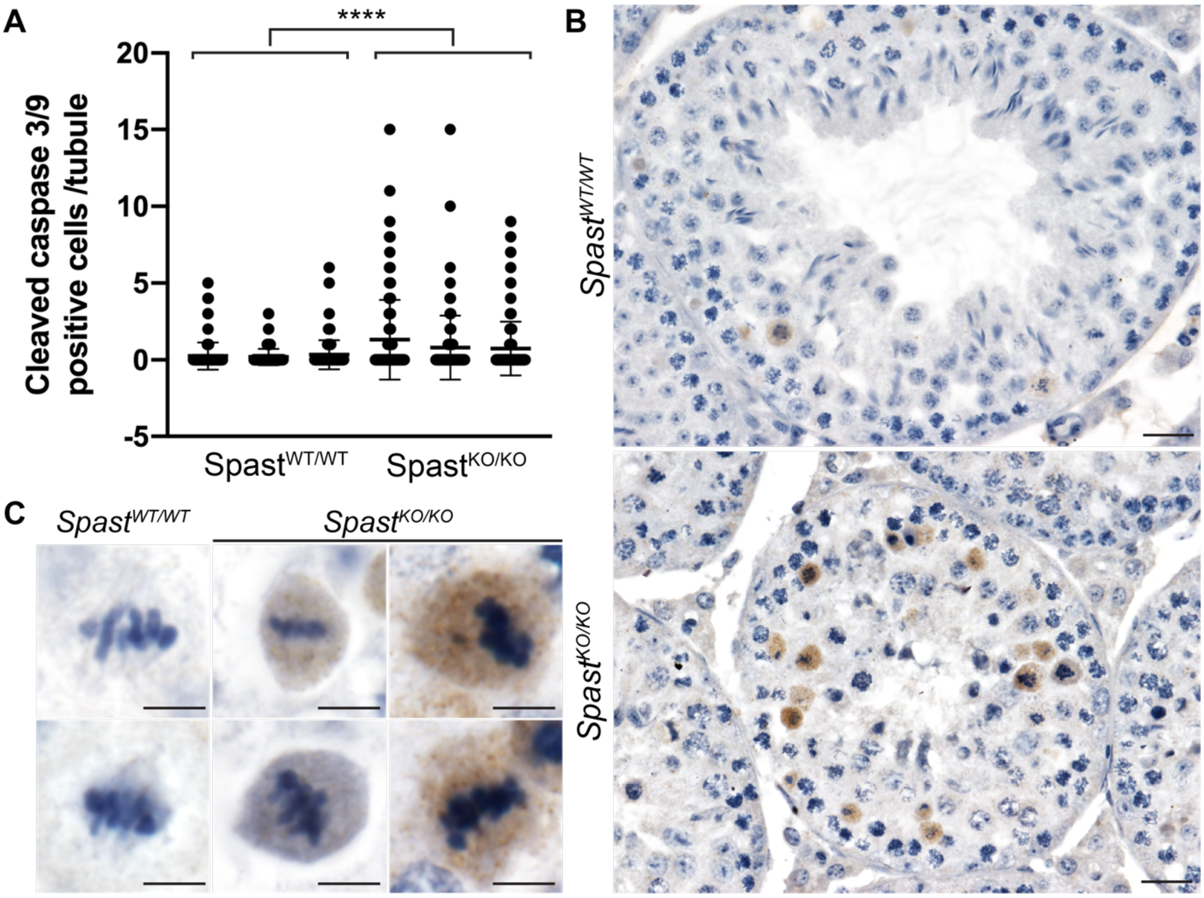


**Fig. S1: Loss of spastin results in an increase of germ cell apoptosis**. Apoptosis of germ cells was assessed using immunohistochemical staining of cleaved-caspase 3 and 9. The average number of cleaved-caspase 3 and/or 9 positive cells per seminiferous tubule per mouse is graphed in (A). Each column represents a single mouse and lines represent mean ± s.d. A minimum of 100 randomly selected seminiferous tubules per mouse were counted. A statistically significant increase in germ cell apoptosis was found in *Spast*$^{KO/KO}$ mice compared to *SpastWT/WT* mice, **** $p<0.0001$. Representative seminiferous tubules and cells for each genotype can be seen in (B-C). Scale bars in B = 20µm, scale bars in C = 5µm.

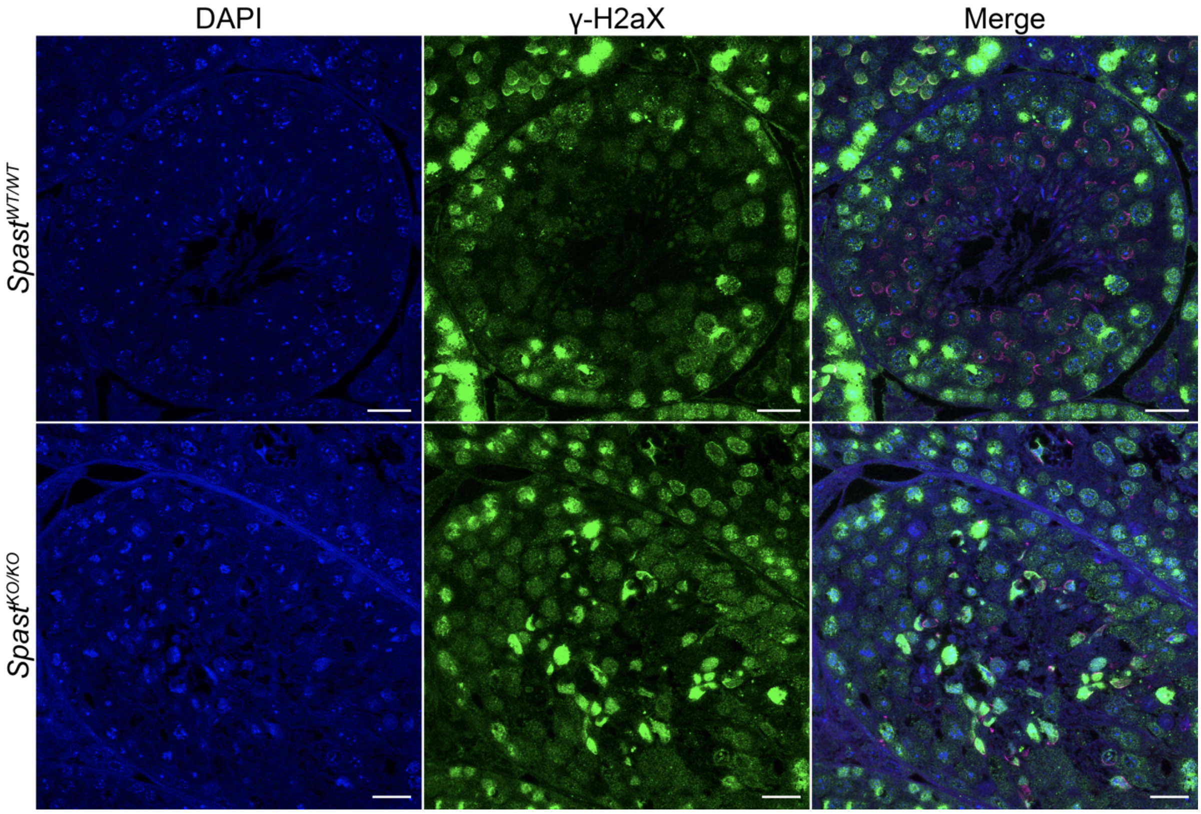


**Fig. S2: The loss of spastin results in an increase in DNA double stranded breaks in post-meiotic germ cells.** Staining for γ-H2aX (green) to identify double stranded breaks in DNA identified staining in post-meiotic spermatids from *Spast*$^{KO/KO}$, but not *Spast*$^{WT/WT}$ mice. Nuclei are counterstained with DAPI (blue). Post-meiotic spermatids may be identified by their location closer to the tubule lumen and by the presence of an acrosome, counterstained with PNA (magenta) in the merged image. Scale bar = 20μm.
